# A Comprehensive Benchmark of Tools for Efficient Genomic Interval Querying

**DOI:** 10.1101/2025.02.19.639167

**Authors:** Richard A. Schäfer, Rendong Yang

**Affiliations:** Department of Urology, Northwestern University Feinberg School of Medicine, Chicago, IL 60611; Department of Urology & Robert H. Lurie Comprehensive Cancer Center, Northwestern University Feinberg School of Medicine, Chicago, IL 60611

**Keywords:** interval, query, data, genomic features

## Abstract

Efficiently querying genomic intervals is fundamental to modern bioinformatics, enabling researchers to extract and analyze specific regions from large genomic datasets. While various tools have been developed for this purpose, there lacks a comprehensive comparison of their performance, memory usage, and practical utility. We present a systematic evaluation of genomic interval query tools using simulated datasets of varying sizes. Our benchmarking framework, `segmeter`, assesses both basic and complex interval queries, examining runtime performance, memory efficiency, and query precision across different tools and data structures. This comprehensive analysis provides insights into the strengths and limitations of different approaches to genomic interval querying, offering guidance for tool selection based on specific use cases and data requirements. The `segmeter` framework and all benchmark data are freely available, facilitating reproducibility and enabling researchers to conduct their own comparative analyses.

## 1 Introduction

Efficiently querying specific genomic regions is fundamental in bioinformatics, enabling researchers to extract relevant information from large genomic datasets. Genomic data files often contain billions of records, making retrieving data without specialized tools computationally expensive. Querying by genomic regions allows researchers to focus on specific regions of interest, such as protein-coding regions to characterize the consequences of genetic variants on the proteome [1, 2]. In addition, genomic queries are integral to numerous algorithms that assign biological data to genomic feature elements. For example, querying genomic regions is crucial in peak calling to identify enriched regions linked to functional elements [3], in homology detection to find conserved sequences across species [4, 5, 6], and in RNA-RNA interaction prediction to pinpoint binding sites within specific gene regions, such as UTRs [7]. Tools for querying specific genomic intervals are essential for efficiently analyzing large-scale genomic data. To facilitate rapid retrieval of relevant data from large datasets, specialized tools have been developed to allow users to focus on regions of interest. Tools for querying genomic intervals can be broadly categorized based on how they structure data for efficient retrieval. While some methods explicitely generate external index files to accelerate lookups, others dynamically construct in-memory data structures. [8] introduced `tabix`, a tool that applies the concept of indexing to position-sorted and compressed tab-delimited formats. By generating an index, `tabix` enables fast, region-based queries, significantly improving the efficiency of working with large genomic datasets and extending the indexing approach used in alignment data formats like BAM to general intervals. Other tools have been developed to support the querying of genomic regions, such as `BEDtools` [9] and `BEDops` [10]. `BEDtools` is a versatile suite of utilities that allows the manipulation and analysis of genomic intervals in BED, GFF, VCF, and other file formats. It incorporates a hierarchical indexing scheme used in the UCSC genome browser to internally index genome coordinate ranges [11, 12, 13]. In that regard, the UCSC provides command-line utilities for interval data manipulation [14]. `BEDops`, another powerful tool, specializes in optimizing the performance of operations on BED files. It requires the interval data to be sorted, which allows the processing of intervals sequentially rather than loading the interval data into memory, thereby reducing runtime and computational requirements. Similarly, `gia` [15] is designed to stream sorted interval data but also supports operations on unsorted data, by which the data is first loaded into memory. In `bedtk` [16], implicit interval trees are utilized to overlap lists of intervals. Rather than storing explicit pointers between nodes, implicit interval trees maintain a binary search tree using array indices to represent parent-child relationships. This reduces the memory overhead substantially, but preserves efficient interval query capabilities. Based on this concept, the `COITrees` library (https://github.com/dcjones/coitrees) improves the query performance by storing the nodes in a van Emde Boas layout [17]. This library is utilized in `granges` [19], providing genomic interval manipulation capabilities comparable to `BEDTools`. In `GIGGLE` [20], the intervals are stored in a B+ tree to allow large-scale interval comparisons. Other data structures that can be used for querying intervals overlaps include the nested containment list (`NCList`) [22] and the augmented interval list (`AIList`) [18]. In this work, we benchmarked available command-line tools for querying genomic intervals, evaluating their runtime performance and memory requirements, storage and general usability.

## 2 Methods

We systematically compared the performance of various methods for searching interval data (Table 1). For that, we created an integrative benchmarking framework termed `segmeter` that is freely available at https://github.com/ylab-hi/segmeter. Figure 1 provides an overview of the framework and illustrates the modes of operation: data simulation (mode `sim`) and benchmarking (mode `bench`).

**Table 1:**
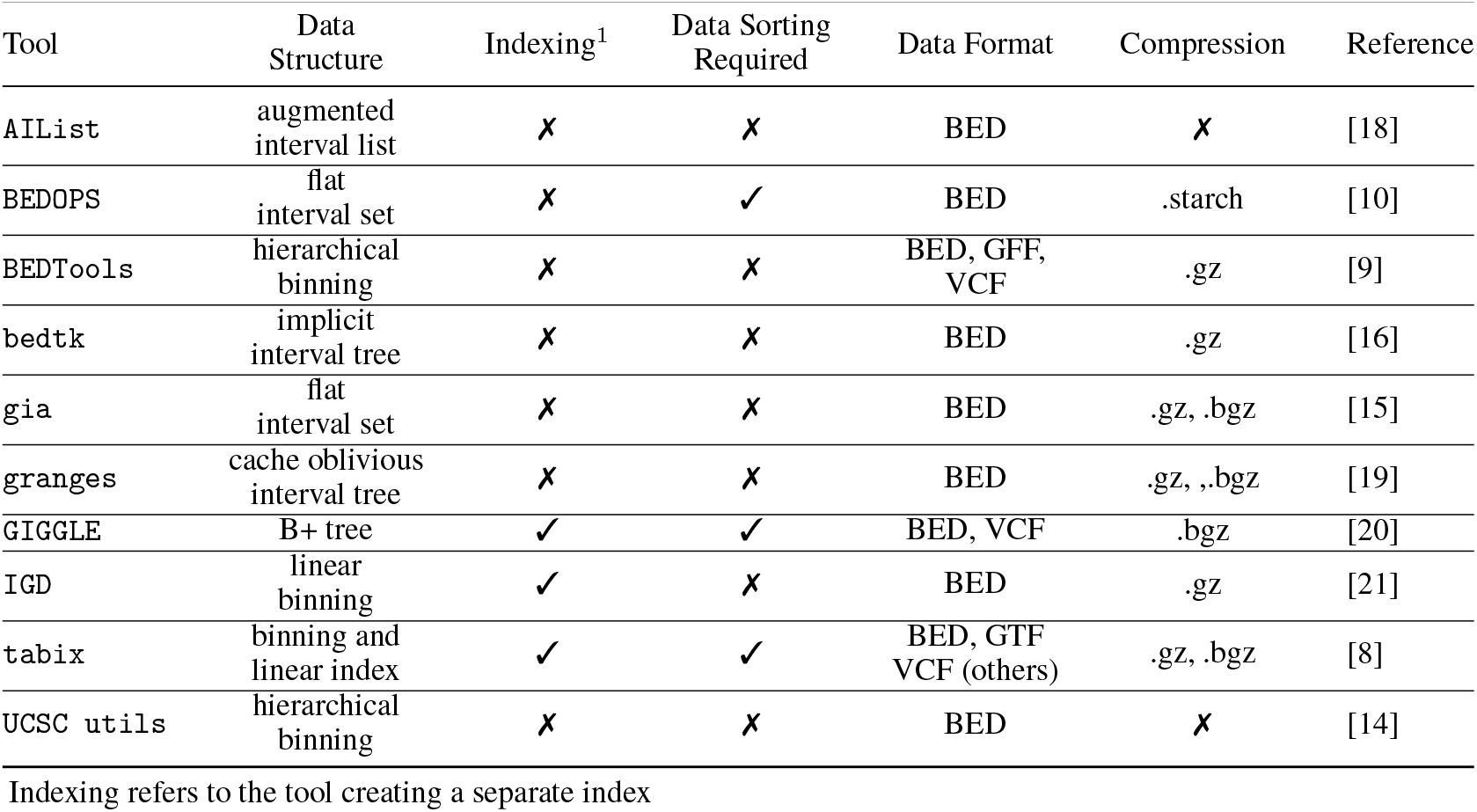
Tools capable for querying genomic intervals.

| Tool | Data Structure | Indexing<sup>1</sup> | Data Sorting Required | Data Format | Compression | Reference |
| --- | --- | --- | --- | --- | --- | --- |
| AiList | augmented interval list | ✗ | ✗ | BED | ✗ | [18] |
| BEDOPS | flat interval set | ✗ | ✓ | BED | .starch | [10] |
| BEDTools | hierarchical binning | ✗ | ✗ | BED, GFF, VCF | .gz | [9] |
| bedtk | implicit interval tree | ✗ | ✗ | BED | .gz | [16] |
| gia | flat interval set | ✗ | ✗ | BED | .gz, .bgz | [15] |
| granges | cache oblivious interval tree | ✗ | ✗ | BED | .gz, .bgz | [19] |
| GIGGLE | B+ tree | ✓ | ✓ | BED, VCF | .bgz | [20] |
| IGD | linear binning | ✓ | ✗ | BED | .gz | [21] |
| tabix | binning and linear index | ✓ | ✓ | BED, GTF VCF (others) | .gz, .bgz | [8] |
| UCSC utils | hierarchical binning | ✗ | ✗ | BED | ✗ | [14] |

Indexing refers to the tool creating a separate index

**Figure 1:**
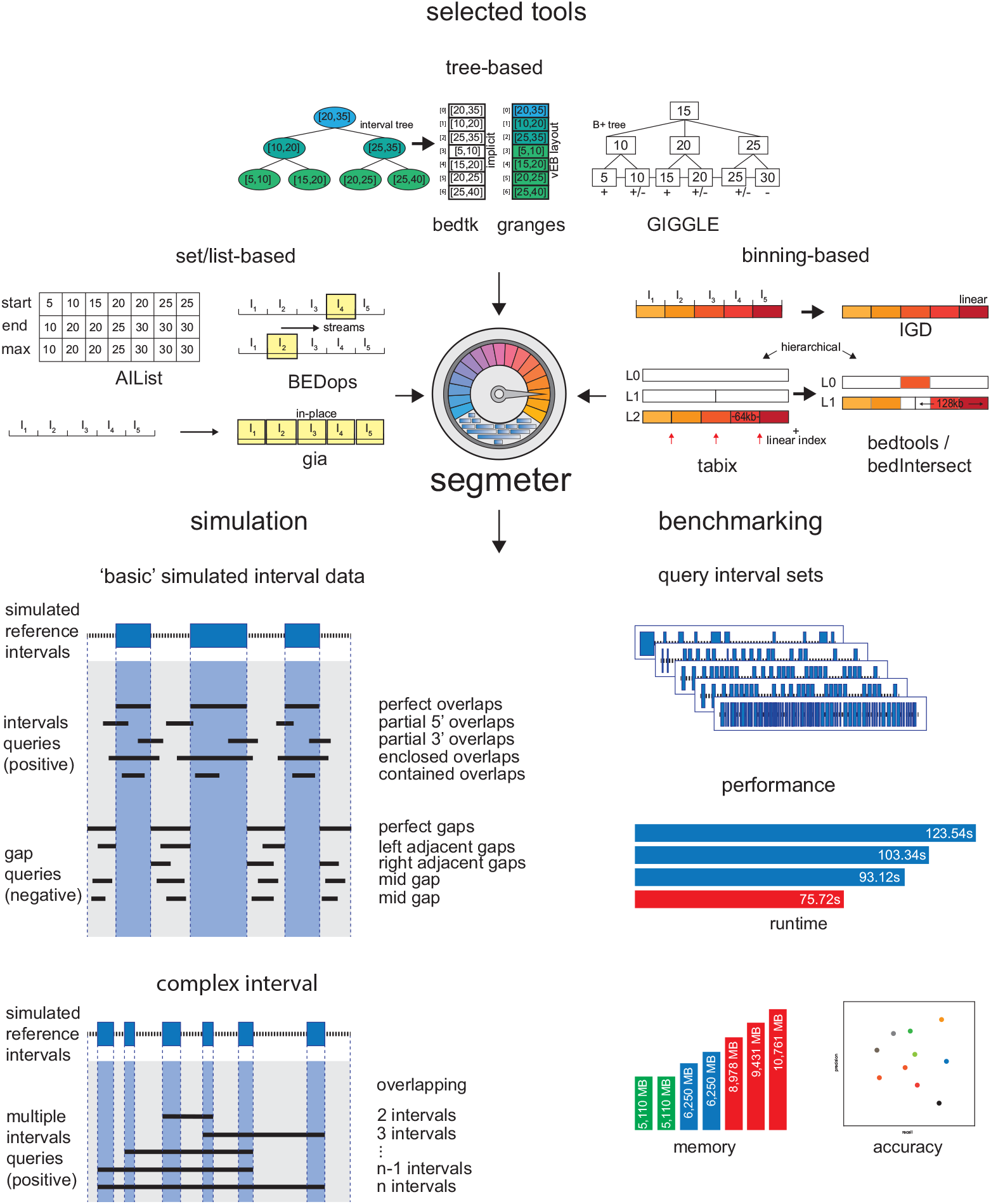
Overview of the `segmeter` framework, the benchmarked methods and the data generated

### 2.1 Data simulation mode

In the data simulation mode, artificial intervals are generated randomly and placed on chromosomes with customizable properties. These include the number of intervals (option --`intvlnums`), the size of each interval (option --`intvlsize`, default: `100-10000`), and the gap size between intervals (option --`gapsize`, default: `100-5000`) which are generated within a user-defined minimum and maximum range. The intervals are placed sequentially on chromosomes, ensuring that most chromosomes have at least ten intervals do not exceed two billion in length. If all chromosomes become fully occupied, any remaining intervals are placed onto additional scaffolds. For each generated reference interval, `segmeter` produces ten corresponding interval and gap queries for benchmarking purposes, as illustrated in Figure 1.

Interval queries are designed to overlap the reference intervals in various ways. These include perfect overlaps, where the query interval is identical to the reference interval, partial overlaps at either the 5’ or 3’ ends, enclosing intervals that fully contain the reference interval, and contained intervals entirely within the reference interval. Gap queries, on the other hand, target the spaces between reference intervals and do not overlap the intervals themselves. These include perfect gaps, where the query matches the gap between two intervals exactly, gaps adjacent to the start or end of an interval, and two gaps entirely contained within larger gaps. In addition, `segmeter` subsamples the resulting query datasets by 10%, 20%, …, up to 100% of their original size. Similarly, the generated intervals are used for complex queries that cover multiple intervals. In particular, `segmeter` generates for *n* intervals on each chromosome queries of different lengths, covering 2 to *n* intervals. These are distributed into bins corresponding to deciles.

### 2.2 Benchmarking mode

In the benchmarking mode, `segmeter` evaluates each tool by assessing the runtime for indexing (if required) and querying the intervals, memory requirements, and precision. Since the methods vary in functionality, we focused the benchmark specifically on detecting overlaps. In particular, the tools are supposed to return the actual interval where the overlap with the queries occurred. Please refer to the supplementary information for more detailed information about how the different tools were executed. In `segmeter`, we included `tabix`, `bedtools` on unsorted interval data, `bedtools` on sorted interval data, `bedtools` using `tabix` for random access, `bedops` and `bedmap` from the `BEDOPS` utility, `giggle`, `gia` on unsorted and sorted data, `bedtk` on unsorted and sorted data, `IGD`, `AIList`, and `bedIntersect` from the UCSC utilities. Benchmarking was primarily performed on unsorted data. For tools requiring preprocessing steps, such as sorting or indexing of reference and query data, we included these computational costs in the indexing and querying runtime, respectively. Precision was determined based on the accuracy of returned overlaps. For basic interval queries, we assessed precision by verifying that the reported overlaps matched the reference intervals. For complex queries, we measured precision by calculating the distance in the number of overlapping intervals. In this mode, `segmeter` generates output files containing statistics for each benchmarked tool (option `--tool`).

### 2.3 System environment

All benchmarks were conducted on a MacBook Pro (2023) with MacOS 15.2, equipped with an Apple M2 chip and 32GB of unified memory. We used the Docker containers provided by `segmeter` the considered tools, which were restricted to a single CPU core and allocated a maximum of 12GB of memory to ensure a fair and reproducible evaluation.

## 3 Results

We used `segmeter v0.12.x` to generate datasets with 10, 100, 1, 000, 10, 000, and 100, 000 intervals. For that, we used the default values for `--intvlsize` and `--gapsize`. This results in 10 to 4287 intervals distributed on 24 chromosomes, with an average length of 5490 for each interval and 2591 for the gaps (see supplementary Table S1, Figure S1, and Figure S2). Subsequently, we evaluated each tool using `segmeter` in three different runs. The resulting metrics were evaluated in the following.

### 3.1 Basic queries

We analyzed the runtime needed for interval and gap queries, with results shown in Figure 2. Additionally, we measured the runtime of `bedtools` and `bedtk` on sorted data, `bedtools` combined with `tabix` for random access, and `bedmap` from the `BEDops` utility. However, as we account for sorting time, we did not observe any runtime improvements over unsorted data (see Figure S3, Figure S4, and Figure S5). For overlapping interval queries (Figure 2A-E), `bedtk` and `AIList` consistently demonstrated superior performance across all interval query sets, with `granges`, `IGD`, and `gia` exhibiting an increased runtime. `tabix` performed adequately for smaller query sets (up to 1,000 queries) but showed poor scaling with increased query volume. While `GIGGLE` and `UCSC utils` showed comparable performance for smaller query sets, `GIGGLE`’s runtime degraded to match `tabix` at higher query volumes, while `UCSC utils` aligned with `BEDops`. Similarly, `bedtools` initially performs comparable to `granges` and `IGD`, but its runtime eventually matches that of `UCSC utils` and `BEDops`. In contrast, `UCSC utils` maintained linear scaling, performing better than `BEDops` with the number of interval queries increasing to 10^5^. For gap queries (Figure 2F-J), which identify regions without overlap of the reference interval, performance patterns were similar with notable exceptions. `GIGGLE` showed improved performance, matching `BEDops` and `UCSC utils`, while `tabix` maintained its poor performance. In `bedtk` and `granges`, we observed an equivalent performance for small query sets which diverged at higher volumes. The runtime hierarchy remained consistent with `bedtk` leading, followed by `AIList`, while `bedtools`, `granges`, `gia`, and `IGD` continued to exhibit a slower runtime. Analysis of the average fraction of runtime spent on interval and gap queries (Figure 2K) revealed consistent proportions across all tools except `GIGGLE`, where interval queries consumed a notably larger share of total runtime. The total runtime across all query types (Figure 2L) showed `bedtk` maintaining the best performance throughout. For smaller query sets (up to 1,000), `granges` and `AIList` performed similarly as second-best, but at higher volumes `granges` aligned with `gia` and `IGD`. Additionally, `UCSC utils`, `bedtools`, and `BEDops` formed a slower tier, while `GIGGLE` and `tabix` exhibited the highest runtimes overall.

**Figure 2:**
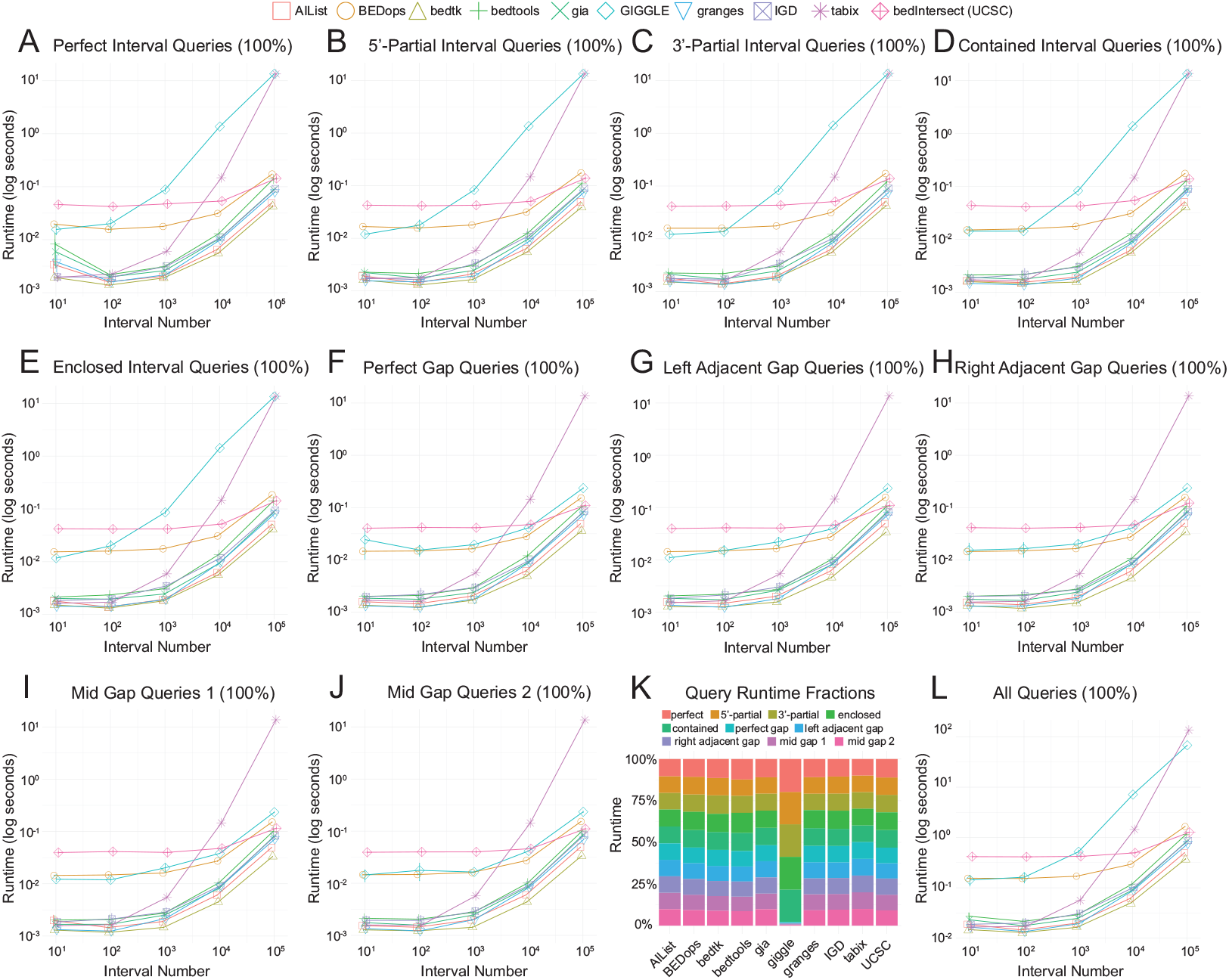
Runtime comparison using simulated interval data. Positive queries **(A-E)** overlap with reference intervals, while negative queries **(F-J)** target gaps between intervals. Panel **K** shows each query’s percentage of total runtime, while panel **L** displays the cumulative runtime.

### 3.2 Complex queries

We analyzed the runtime performance of complex queries, as illustrated in Figure 3. First, we summarized the overall runtime for all complex queries combined (Figure 3A). The results indicate that `AIList`, `bedtk`, and `granges` form a group with consistently fast runtimes across all interval datasets. This is followed by `tabix` and `UCSC utils`, which exhibit moderate runtimes. Another distinct tier, with runtimes approximately ten times higher, includes `gia`, `BEDops`, `IGD`, and `bedtools`. Notably, `GIGGLE` demonstrates the highest runtime, which increases further as the set size of the interval increases. A similar pattern can be observed when inspecting the runtime for each dataset individually (Figure 3B-F). For small datasets (Figures 3B-C), the runtime decreases when the query spans nearly all intervals (10th decile), while the runtime in larger datasets remains constant over different deciles (Figures 3E-F). Interestingly, with an increasing number of reference intervals (Figures 3D-F) we observe a decrease in the runtime per query.

**Figure 3:**
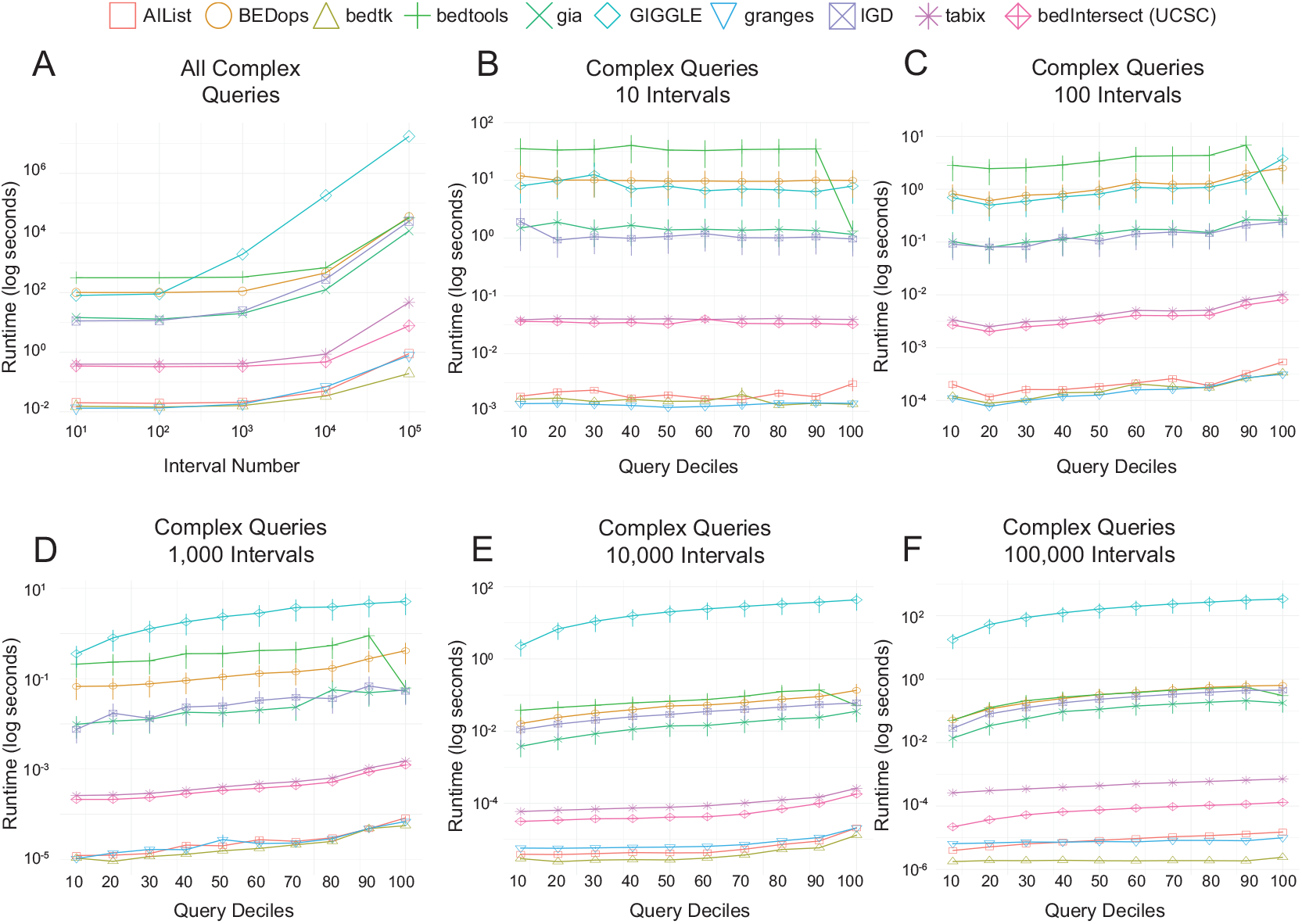
Runtime comparison using simulated complex interval data. Runtime for all complex queries over all interval datasets (A), and averaged runtime per interval query for datasets with 10 (B), 100 (C), 1, 000 (D), 10, 000 (E), and 100, 000 intervals.

### 3.3 Memory requirements

We also analyzed the memory requirements of each tool when querying different interval datasets, as shown in Figure 4. When performing all basic queries (Fig. 4A), `AIList`, `bedtk`, and `IGD` demonstrated the lowest memory usage. A second tier follows, consisting of `tabix`, `UCSC utils`, `GIGGLE`, `BEDops`, and `granges`, which exhibit moderate memory consumption. In contrast, `gia` and `bedtools` require approximately twice the memory of the second tier, placing them in the highest memory usage category. For complex queries, we observed distinct memory usage patterns across tools. `AIList`, `bedtk`, and `IGD` remained the most memory-efficient, while `gia` exhibited reduced memory consumption compared to its usage in basic queries, now aligning with `tabix`, `UCSC utils`, `bedtools`, and `granges`. Meanwhile, `BEDops` showed increase memory usage, which is only exceeded by `GIGGLE`.

**Figure 4:**
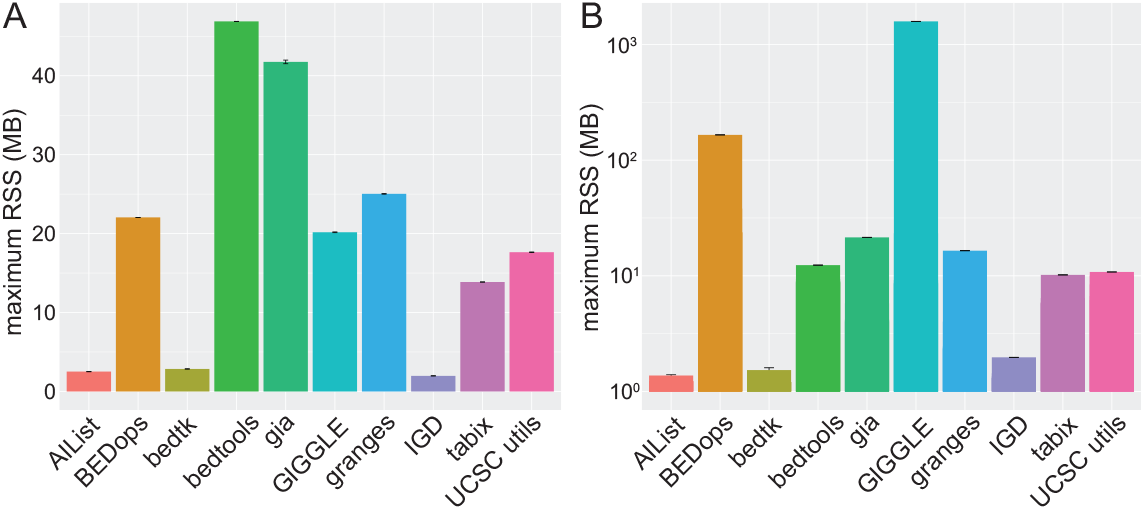
Peak memory usage over all interval datastes for basic (A) and complex queries (B).

### 3.4 Indexed creation

We analyzed the runtime required for indexing various tools, as shown in Figure 5. While indexing is a one-time cost, it contributes to the overall runtime when querying intervals. In the selected tools in this benchmark, only `GIGGLE`, `IGD`, and `tabix` require a preprocessed index when querying interval. Other tools such as `BEDops` simply require the data to be sorted. We included simple sorting and sorting with compression. Our results show that `GIGGLE` exhibits the highest runtime and memory usage across all datasets. In contrast, `IGD` maintains a consistent runtime and memory footprint regardless of the dataset size. It can be seen that `tabix` compared to simple sorting only demonstrate comparable performance, though they experience a significant increase in runtime and memory usage under larger datasets.

**Figure 5:**
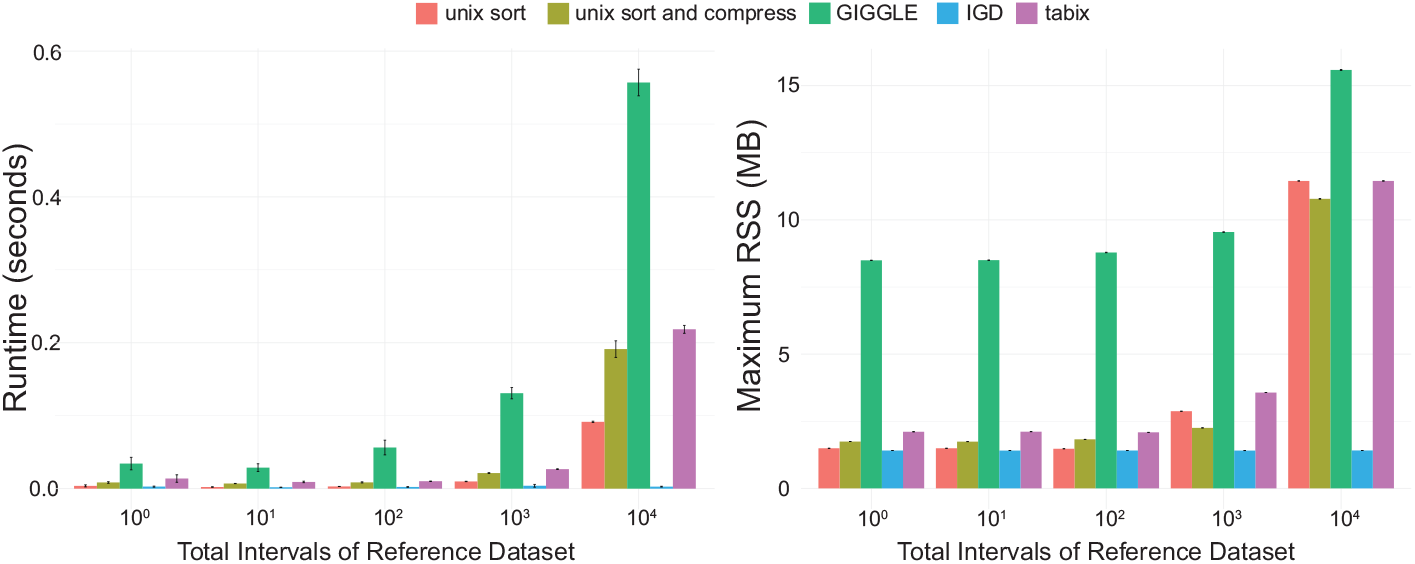
Memory requirements for index creation

### 3.5 Accuracy

When the selected tools are benchmarked, `segmeter` also captures the precision of queries that overlap with reference intervals. We primarily used this for internal purposes to ensure that the tools function correctly and that all overlapping intervals can be retrieved. For that, `segmeter` is configured to process the output of a selection of tools, so that the results are comparable (see supplementary information). With the exception of `granges`, we encountered perfect precision when querying the tools with our generated datasets (see Table S4).

### 3.6 Usability

Most tools could be easily installed using a designated package manager, such as `bedtools` and `tabix` via APT (https://salsa.debian.org/apt-team/apt), `gia` and `granges` via Cargo (https://github.com/rust-lang/cargo), while others, like `AIList`, `BEDops`, `bedtk`, `IGD`, and `GIGGLE` require compilation using Make. We made modifications to the data handling in `GIGGLE`, so it could run correctly (see supplementary information). In most tools, the functionality is restricted to simple overlapping queries, while `BEDops`, `BEDTools`, and `gia` provide a comprehensive feature set.

## 4 Discussion

In this study, we evaluated the performance of various genomic interval query tools using simulated datasets ranging from 10 to 100, 000 intervals, providing insights into the scalability and efficiency of the established methods. Although we observed a relationship between the tools and their data structures, other factors, such as implementation and data handling, may also play a role. This is beyond the scope of this benchmark. Our results showed that the implicit interval tree implementation in bedtk consistently demonstrated superior performance, followed by the augmented interval list approach in `AIList`. The efficiency of these data structures stems from their ability to balance memory efficiency and query speed—implicit interval trees eliminate pointer overhead while preserving efficient interval query capabilities, whereas augmented interval lists achieve fast lookups through strategic augmentation of interval endpoints. In contrast, tools employing more complex data structures, such as the B+ tree in `GIGGLE`, exhibited poorer scalability with larger datasets, suggesting that the overhead of maintaining these structures outweighs their theoretical advantages for genomic interval queries. The B+ tree structure, despite its higher maintenance overhead, can offer advantages in more complex scenarios such as pangenome analysis, in which multiple overlap relationships beed to be handled. For complex queries spanning multiple intervals, we found that the choice of data structure became even more critical. Tools utilizing simpler, memory-efficient structures, such as `bedtk` or `AIList`, exhibited consistent performance across dataset sizes. In contrast, the performance tools with more elaborate indexing schemes diminished. This suggests that for genomic interval queries, the overhead of maintaining complex tree structures may not be justified by the performance benefits, particularly when dealing with large-scale data. Memory usage patterns also reflected the efficiency of different data structures. The low memory footprint of `AIList`, `bedtk`, and `IGD` demonstrates the advantages of their streamlined data representations. In contrast, tools like `bedtools`, which load more data into memory, showed higher memory requirements. This trade-off between memory usage and query performance appears to be a crucial consideration in tool design, with the most successful tools finding an optimal balance. In the indexing requirements, we observed that the B+ structure of `GIGGLE` has the highest indexing costs, while simpler indexing approaches such as in `IGD` demonstrated more practical trade-offs between index creation time and query performance. This suggests that for the considered genomic interval queries, complex indexing strategies may not provide sufficient performance benefits to justify their computational overhead. It remains to be investigated to what extent these methods can demonstrate their usefulness when handling larger interval queries and complex datasets. We found that tools requiring sorted input, such as `BEDops` to show competitive performance, but at the cost of preprocessing overhead. When this proprocessing is accounted for, we found no improvement to methods working on unsorted data. This highlights the importance of considering not just the query algorithm itself, but also the preprocessing requirements when evaluating tool efficiency. The precision analysis revealed that most tools achieved perfect accuracy, with granges being the notable exception. This suggests that while different data structures can significantly impact performance, they generally maintain accuracy in overlap detection. For general interval queries, our results suggest that simpler, memory-efficient data structures like implicit interval trees and augmented interval lists may be more practical than theoretically optimal but complex structures.

## 5 Conclusion

In this work, we introduced `segmeter`, a comprehensive framework for benchmarking tools used in querying interval data. We evaluated widely used methods in the field, assessing their runtime, memory usage, and overall applicability. While a variety of data structures exist for genomic interval queries, our results suggest that simpler, memory-efficient structures often outperform more complex ones in practical applications. Among the tools tested, `bedtk` and `AIList` demonstrated the highest efficiency on large datasets, whereas others, such as `GIGGLE` and `tabix`, faced scalability challenges. This benchmark primarily assessed the tools themselves; however, their performance is also influenced by the characteristics of the interval data and the implementation details of the underlying data structure.

## 6 Data availability

We have established a permanent data repository on Zenodo (DOI: 10.5281/zenodo.14880992), which contains the simulated data used in our benchmarking. We provide detailed statistics on the simulation data in Table S1. Table S2 contains all runtime measurements and memory usage for the different benchmark runs, while Table S3 presents the same metrics for the index creation process. Finally, Table S4 summarizes the precision results of the benchmarks. The repository includes all relevant scripts and configurations to reproduce our results.

## 7 Code availability

We distribute `segmeter` under the MIT license and make it freely available through GitHub (https://github.com/ylab-hi/segmeter). The repository includes comprehensive documentation, installation instructions, and Docker containers to ensure reproducibility. All benchmarking scripts and configuration files used in this study are also included in the repository.

## 8 Competing interests

RY has served as an advisor/consultant for Tempus AI, Inc. This relationship is unrelated to and did not influence the research presented in this study.

## 9 Author contributions statement

RAS and RY designed the study. RAS conducted the experiments and analysed the results. R.A.S and R.Y wrote and reviewed the manuscript.

## 10 acknowledgments

This project is supported in part by NIH grants R35GM142441 and R01CA259388 awarded to RY.

## Notes

https://zenodo.org/records/14880992?token=eyJhbGciOiJIUzUxMiJ9.eyJpZCI6ImI1OTdlZmFkLWY1YWEtNGFmOS1hNTEwLTdiMTEwYjRiMDg0NyIsImRhdGEiOnt9LCJyYW5kb20iOiIzNGFmOGY1Mzk4NTI5M2NjZWJjNmM4NTc1YmYwNjljZiJ9.wowrt0SMac-VZ79xRZfvc_utfNifnf6A38-AynvZFrgjH0fRaCyiPjXAapYaTmGGnXsYVf1l5TC0DQLaBUQi4w

### Summary of Updates

This is a small update that resolve a typo in the abstract in which 'semester' was written instead of 'segmeter', and the mail address of the corresponding other was wrong.

