## supplementary information for "A Comprehensive Benchmark of Tools for Efficient Genomic Interval Querying"

PAPER

### Execution of external tools

The tools and their corresponding commands are listed below, as they were utilized in `segmeter` (using the --tool option).

#### AIList

`AIList v0.1.1` is utilized and applied on unsorted data using the following command:

```
ailist <reference.bed> <query.bed>
```

However, the output is de-duplicated for overlapping intervals in complex queries. In the fourth column, the number of overlaps is reported, which is then used by `segmeter` to ensure that each overlap is represented individually.

#### BEDops

`BEDops v2.4.41` is utilized, which requires sorted input. Here, we use different commands for basic and complex queries.

```
sort -k1,1 -k2,2n -k3,3n <query.bed> > <query.sorted.bed> # sort the data
bedops --element-of 1 <reference.sorted.bed> <query.sorted.bed> # query basic intervals
bedmap --echo-map --multidelim '\n' <query.sorted.bed> <reference.sorted.bed> # query complex intervals
```

It is to be noted that in `segmeter`, internally the command `bedops` uses the above configuration, while `bedmaps` uses `bedmap` for both basic and complex queries.

#### bedtools

`bedtools v2.30.0` is utilized, and we applied it to unsorted, sorted, and indexed data:

**bedtools (unsorted):** Intersection without preprocessing:

```
bedtools intersect -wa -a <reference.bed> -b <query.bed>
```

**bedtools with sorted data:** Requires pre-sorting of both files:

```
sort -k1,1 -k2,2n -k3,3n <query.bed> > <query.sorted.bed>
bedtools intersect -wa -a <reference.sorted.bed> -b <query.sorted.bed>
```

**bedtools with tabix:** Uses sorted and indexed data:

```
sort -k1,1 -k2,2n -k3,3n <query.bed> > <query.sorted.bed>
bedtools intersect -wa -a <reference.sorted.bed> -b <query.bed>
```

#### bedtk

`bedtk` is utilized and applied to both sorted and unsorted. However, `bedtk` has no official release. Hence, we used the version downloaded on 2025-01-25. We used the following command to query basic and complex interval queries:

```
bedtk flt <query.bed> <reference.bed>
```

© The Author 2022. Published by Oxford University Press. All rights reserved. For permissions, please

However, `bedtk` only reports direct intersections and does not report the original reference intervals (as required for `segmeter`). For that reason, we subsequently applied `bedtools`, which allows us to report the reference intervals from the output of `bedtk`:

```
bedtools intersect -wa -a <filtered.bed> -b <query.bed>
```

However, this was not accounted for in the runtime and memory measurements.

### gia

We utilized `gia v0.2.23` and applied it to unsorted data:

```
gia intersect -a <query.bed> -b <reference.bed> -t
```

### giggle

We encountered several challenges while working with `giggle v0.6.3`:

1. Compilation issues on Ubuntu $\geq$ 20.04 which has been documented in Issue 9 (https://github.com/ryanlayer/giggle/issues/69)
2. The `giggle index` subcall failed to recognize input and output folders sharing the same parent directory

We created a Docker container (yanglabinfo/segmeter:giggle-latest) based on Ubuntu 20.04 (1), forked `giggle` and modified its data handling implementation (fork available at https://github.com/riasc/giggle). In the working implementation, `segmeter` uses internally the following commands:

```
giggle index -i <indexbedfile> -o <indexfolder> -f -s # create the index
giggle search -i <indexfolder> -q <querybedfile> -v # search for overlap against the index
```

### granges

We utilized `granges`, which requires a `--genome` file with the length of each chromosome:

```
granges filter --genome <chromlens_file> --left <reference.bed> --right <query.bed>
```

### IGD

We utilized `IGD v0.1.1` and applied it to unsorted data using the following commands:

```
igd create <input_dir> <output_dir> <label> # create index
igd search <index_dir>/<label>.igd -q <query.bed> -f # query intervals
```

However, the output of `IGD` is not in valid BED format and instead reports the overlaps for each query. For that reason, `segmeter` processes the output and converts it into valid BED format. Again, we did account for this in the runtime and memory measurements.

### tabix

We utilized `tabix v1.16`. The data needs to be sorted and indexed which was done using the following commands:

```
sort -k1,1 -k2,2n -k3,3n <input.bed> > <sorted.bed> # sort the reference data
bgzip -f <sorted.bed> > <sorted.bed.gz> # compress the data
tabix -f -C -p bed <sorted.bed.gz> # index the data
```

Subsequently, we can query the index reference data using:

```
tabix <sorted.bed.gz> -R <query.bed>
```

### UCSC Utils

We used `UCSC utils` from the UCSC utilities (http://hgdownload.soe.ucsc.edu/admin/exe/) and was used in the following command:

```
bedIntersect -aHitAny <input.bed> <query.bed>
```

Similarly, to `bedtk`, we used `bedtools` to report all overlaps.

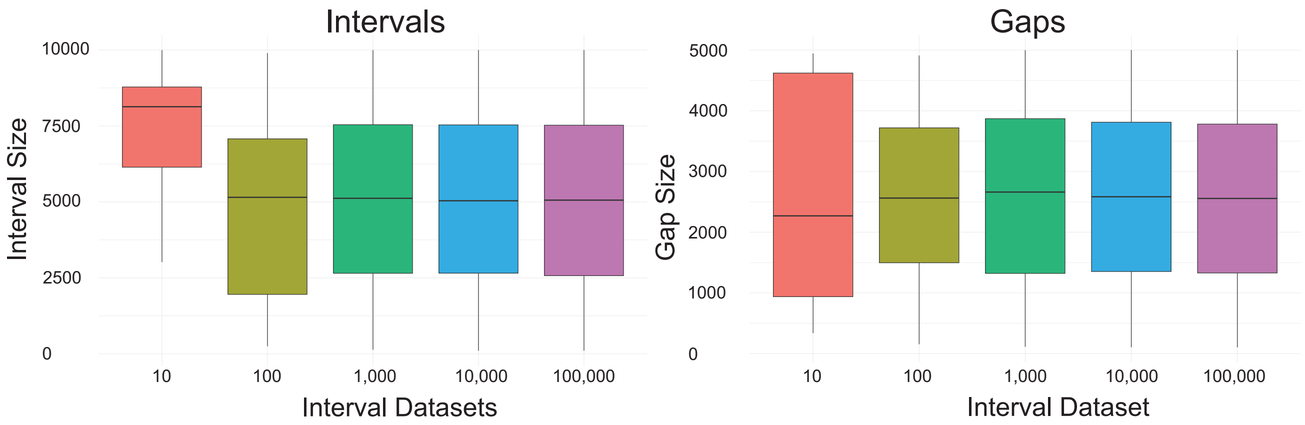


**Fig. 1.** Interval and gap sizes of the generated interval data.

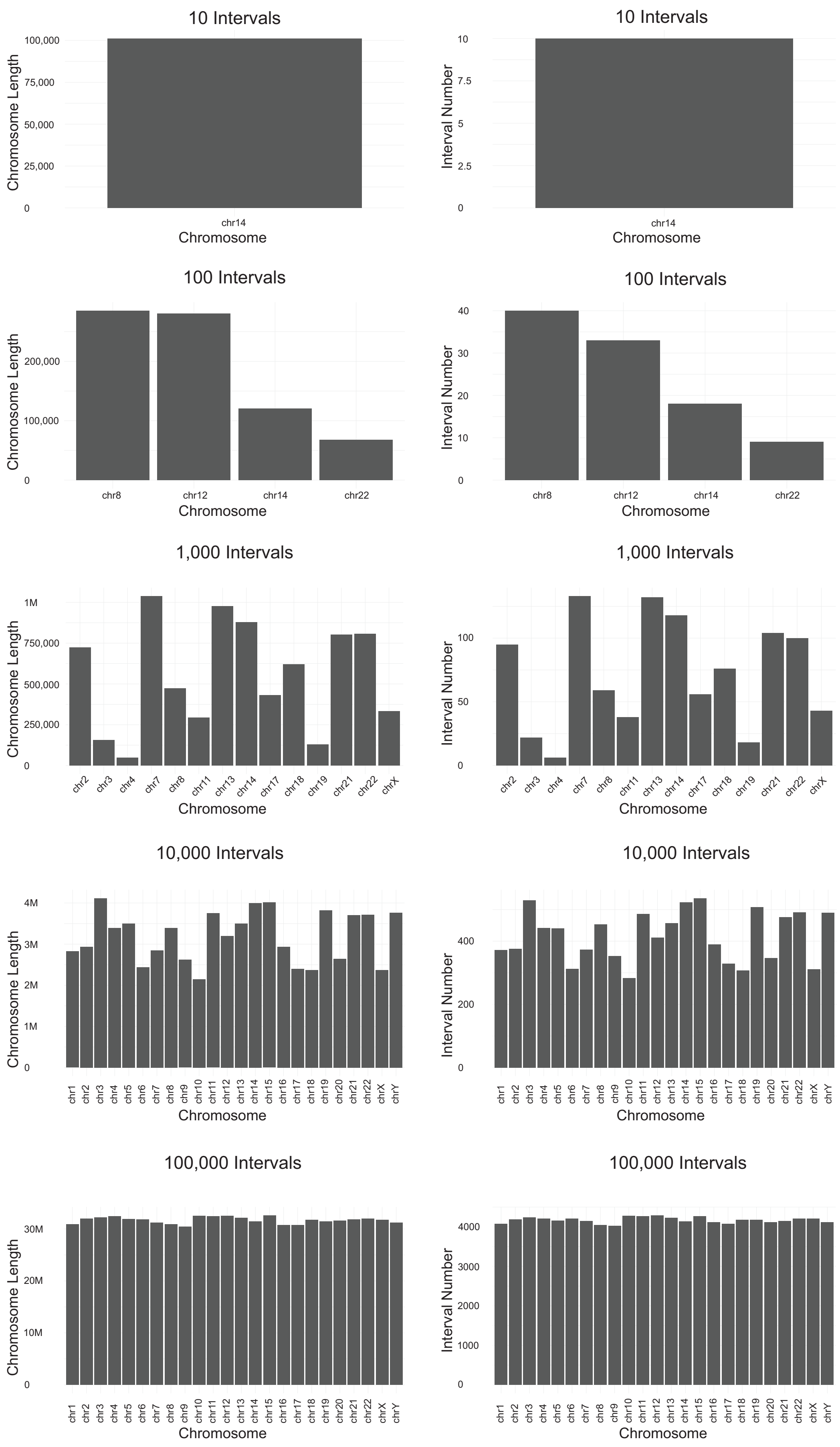


**Fig. 2.** Chromosome length and interval per chromosome distribution of the simulated datasets

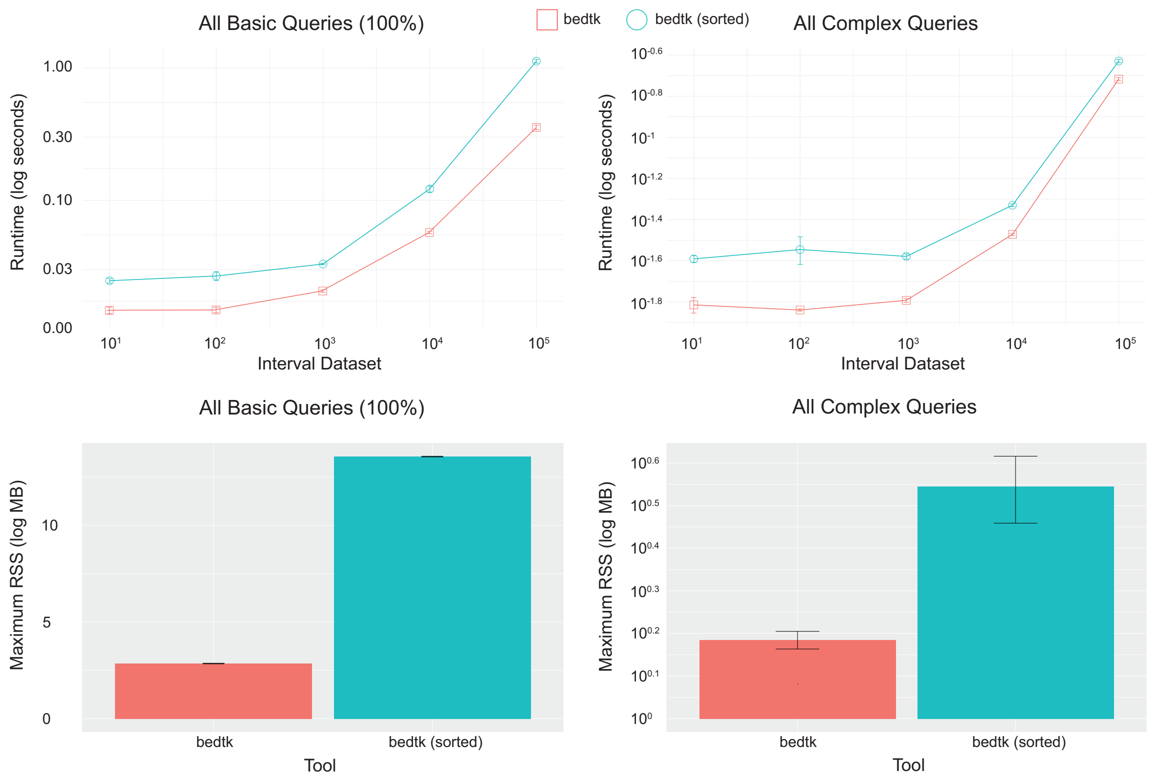


**Fig. 3.** Runtime memory requirements for `bedtk` compared to `bedtk` on sorted data

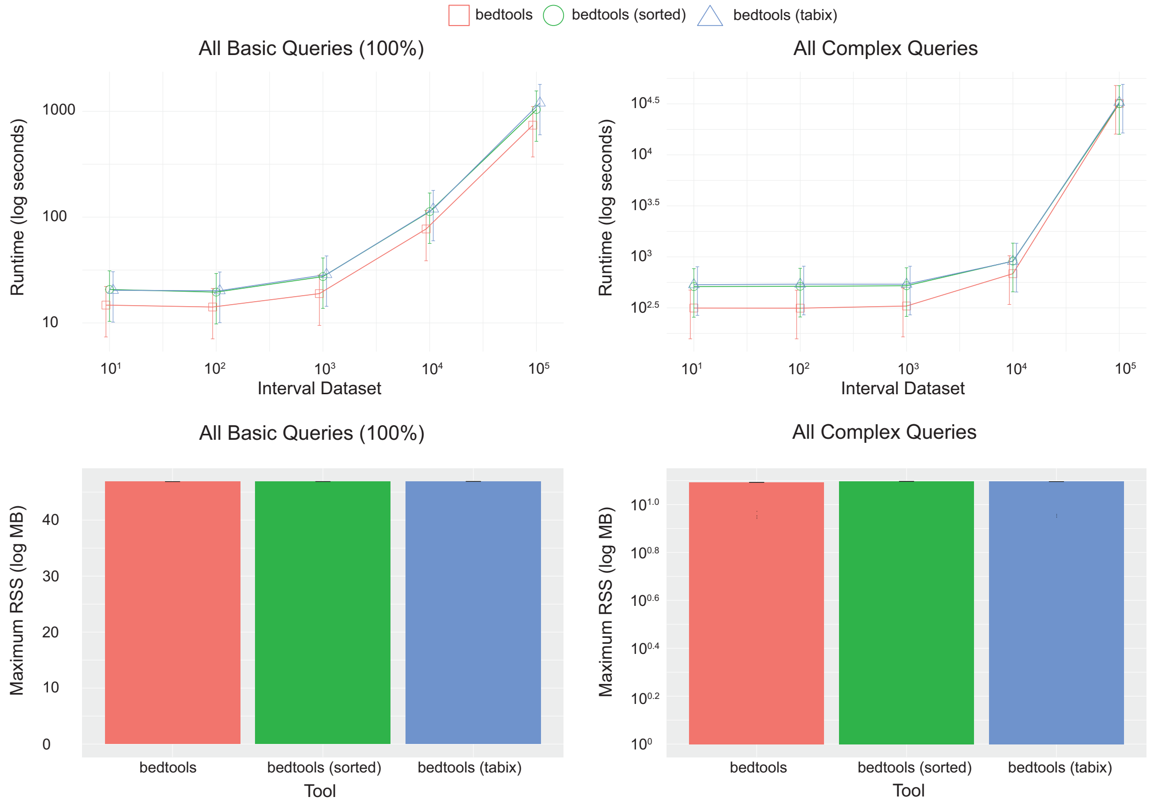


**Fig. 4.** Runtime and memory requirements for `bedtools` compared to sorted data, and with random access by `tabix`

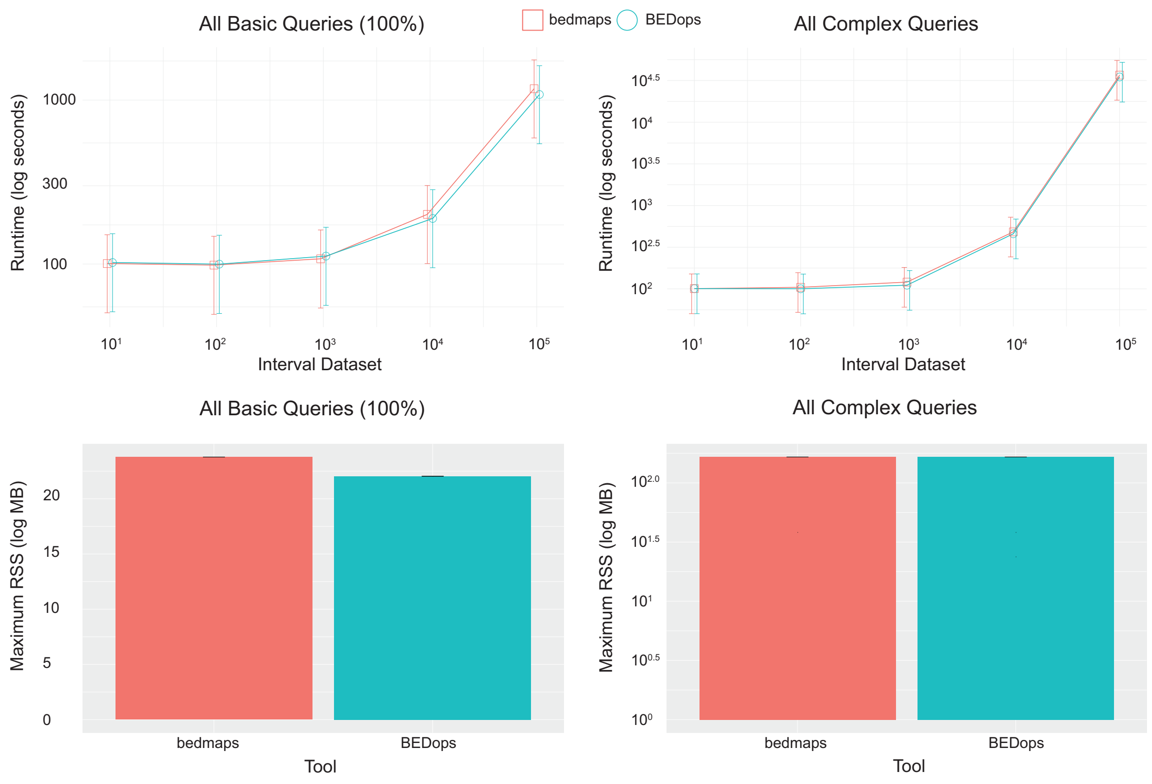


**Fig. 5.** Comparison of the runtime and memory requirements for `BEDops` and `bedmap` from the `BEDops` utilities in basic and complex queries
